# Transcriptional activation by MNRR1 is effected by recruiting p300 and can be induced by minimal peptides

**DOI:** 10.1101/2025.08.25.672219

**Authors:** Neeraja Purandare, Vignesh Pasupathi, Deepesh Padhan, Sagarika Rai, Lawrence I. Grossman, Siddhesh Aras

**Author notes:** Current position, University of Toledo (Neeraja Purandare).

## Abstract

Mitochondrial Nuclear Retrograde Regulator 1 (MNRR1; also, CHCHD2, PARK22, AAG10), which functions in both the mitochondria and the nucleus, modulates mitochondrial function as well as cellular stress response. We have previously shown that stress response is predominantly mediated by its nuclear function as a transcriptional regulator; it operates at a conserved 8-bp DNA element that responds to moderate hypoxia, that we have termed the oxygen responsive element (ORE). This 8-bp element is the consensus DNA binding site for the transcription factor Recombination Signal Binding Protein For Immunoglobulin Kappa J Region (RBPJκ). We previously addressed the mechanism of nuclear MNRR1 by showing that it displaces the inhibitory transcriptional regulator CXXC-type zinc finger protein 5 (CXXC5), facilitating activation of genes harboring the ORE, and that it most effectively does so after deacetylation by SIRT1 at Lys119. A deacetyl mimetic mutant (K-R) rescues the phenotype in both genetic and environmental inflammatory models of defective mitochondrial function. Here we have refined the mechanism by which MNRR1 regulates transcription at the ORE. We show that MNRR1 interacts with RBPJκ and recruits the transcriptional co-activator p300 to facilitate transcription. We also show that a minimal domain of MNRR1 is sufficient to activate its nuclear function. Peptides based on this minimal domain can activate transcription by MNRR1 by enhancing p300 and RBPJκ interaction. MNRR1 peptides activate downstream pathways such as mitochondrial biogenesis and the unfolded protein response (UPR^mt^) in an *in vitro* model for MELAS.

## Introduction

MNRR1 (also known as CHCHD2, PARK22, AAG10) is a biorganellar regulator of mitochondrial function that works by acting in two compartments, the nucleus and the mitochondria. MNRR1 is an important stress-responsive gene that is associated with a number of physiological and pathological conditions including neurodegeneration [1–6], inflammation [7–10], and cancer [11, 12]. At the sub-cellular level, MNRR1 functions in a distinct manner in each compartment. Nuclear MNRR1 acts as a transcriptional regulator [13–15] that can enhance the transcription [13] of several genes associated with mitochondrial function and stress response [1, 16]). Mitochondrial MNRR1 has two distinct known roles: a) it can directly bind cytochrome *c* oxidase (complex IV) of the electron transport chain and regulate oxygen consumption and ATP production [2, 16, 17], and b) it can bind Bcl-xL to regulate apoptosis [18].

Previously, we showed that under moderate hypoxia, MNRR1 along with RBPJ_κ_, act together to control the transcription of *COX4I2* (cytochrome *c* oxidase subunit 4 isoform 2) that harbors an 8-bp DNA responsive element in the promoter termed the oxygen responsive element (ORE). The same element is also present on the promoter of *MNRR1* itself [16], suggesting that MNRR1 can regulate its own transcription [1, 16]). We have recently identified that under pseudohypoxia, as seen in conditions such as MELAS (mitochondrial encephalomyopathy, lactic acidosis and stroke-like episodes), further regulation occurs via HIF2α [19]. A HIF2α binding element (hypoxia responsive element or HRE) overlaps the ORE and can inhibit RBPJ_κ_ and MNRR1-mediated transcriptional activation of *MNRR1,* resulting in reduced levels of MNRR1 protein and thereby contributing to the mitochondrial deficits seen in MELAS [19].

Here we have further characterized the role of MNRR1 in transcriptional regulation via RBPJκ and defined a minimal MNRR1 peptide that can carry out transcriptional activation. The mechanism by which RBPJ_κ_ is activated by the Notch signaling pathway [20–22] has been well established and consists of inhibitory proteins such as SHARP (SMRT/HDAC1 Associated Repressor Protein) [23], which recruits KMT2D (Lysine Methyltransferase 2D). Upon activation of this signaling pathway, the histone acetyltransferase p300 is recruited to activate transcription [24]. We now identify the role of MNRR1 in this complex, which promotes activation of RBPJ_κ_ mediated transcription. We further perform a functional analysis by serially deleting 15 amino acid segments of the full length MNRR1 protein to identify the minimal region of the protein required for this nuclear function. Since MNRR1 activation, specifically the nuclear function, is sufficient to rescue deficits associated with MELAS [1], and also placental inflammation [8], we propose to use this as a novel method to enhance MNRR1 nuclear function in pathogenic conditions where these levels are reduced. Peptide drugs have been in use for over a decade since the discovery of insulin and, more recently, GLP-1 agonists [25]. To date, over 170 peptide drugs have been identified due to various advantages including high specificity and efficacy and low immunogenicity [26]. Hence, identifying a peptide drug that mimics the transcriptional activation function of MNRR1 in such conditions could be a novel therapeutic avenue to target inflammatory and neurodegenerative conditions where MNRR1 is downregulated.

## Materials and Methods

### Cell Lines

The 143B cells containing 73% of the MELAS m.3243A>G mtDNA mutation (DW7, MELAS cells), CL9 cells (100% WT mtDNA), and the NARP cells were a kind gift from Dr. Douglas Wallace (Children’s Hospital of Philadelphia, PA) and were grown in DMEM with 1 mM pyruvate supplemented with non-essential amino acids (Gibco, Billings, MO, USA), 50 μg/mL uridine (Sigma Aldrich, Burlington, MA, USA), and 10% FBS (Fetal Bovine Serum, Sigma Aldrich, Burlington, MA, USA) plus penicillin-streptomycin. The Rho^0^ cells (143B background) were a kind gift from Dr. Eric Schon (Columbia University, NY) and the MELAS patient fibroblasts were a kind gift from Dr. Tamas Kozicz (Icahn School of Medicine at Mount Sinai, NY). The patient fibroblast and Rho^0^ cells were cultured in DMEM with 1 mM pyruvate supplemented 50 μg/mL uridine (Sigma Aldrich, Burlington, MA, USA), and 10% FBS (Sigma Aldrich, Burlington, MA, USA) plus penicillin-streptomycin. The human first trimester placental cells HTR8/SVNeo (HTR) were obtained from the ATCC (Manassas, VA, USA) and were cultured in Roswell Park Memorial Institute Medium (RPMI) (HyClone, Logan, UT, USA), supplemented with 5% FBS plus penicillin-streptomycin. The primary retinal endothelial cells were a kind gift from Dr. Jena Steinle (Wayne State University, Detroit, MI) and were grown in Cell Systems normal glucose (5mM) medium (Kirkland, WA, USA) supplemented with microvascular growth factors (MVGS), 10ug/mL gentamycin, and 0.25ug/mL amphotericin B (Invitrogen, Carlsbad, CA, USA) on attachment factor coated dishes. The LHON mutant LCLs (Lymphoblastoid cell line, 3460G>A mtDNA mutation) were obtained from Coriell (Camden, New Jersey, USA) were cultured in RPMI (HyClone, Logan, UT, USA), supplemented with 15% FBS plus penicillin-streptomycin. The NPC1 (Niemann-Pick C1) mutant LCLs (Coriell, Camden, New Jersey, USA) were also grown in RPMI (HyClone, Logan, UT, USA), supplemented with 20% FBS plus penicillin-streptomycin. The MNRR1-KO (MDA-MB-231) cells were grown in DMEM with 1 mM pyruvate supplemented with and 10% FBS (Sigma Aldrich, Burlington, MA, USA) plus penicillin-streptomycin and puromycin (1 μg/mL, Invivogen, San Diego, CA, USA).

### Chemicals

All versions of the MNRR1 peptide (full length petide, scrambled peptide, and FITC-conjugated versions as well as peptide deletions were purchased from Peptide 2.0 (Chantilly, Virginia, USA) and solubilized in cell-culture grade water. LPS (Lipopolysaccharide from Escherichia coli 0111:B4) was purchased from Invivogen (San Diego, CA, USA). Peptide at the indicated concentrations were added to the cell culture media and incubated for 48 hrs prior to downstream analysis.

### Plasmids

The MNRR1 1kB promoter luciferase reporter plasmid and pRL-SV40 *Renilla* luciferase expression plasmids and the full length MNRR1-expressing plasmid have been described previously [13]. The promoterless luciferase reporter plasmid was obtained from Promega (Madison, WI, USA). The MNRR1 promoter luciferase reporter with a deletion of the ORE has been described previously [19]. The plasmids for expression of MNRR1 protein deletions of 15 amino acids were generated using overlap extenstion [16]. All expression plasmids were purified using the EndoFree plasmid purification kit from Qiagen (Germantown, MD, USA).

### Transient Transfection

MELAS cells were transfected with the indicated plasmids using ViaFect transfection reagent (Promega, Madison, WI, USA) according to the manufacturer’s protocol. A reagent–DNA ratio of 3:1 in complete medium was used. Following incubation at room temperature for ∼15 min, the cells were overlaid with the mixture. The plates were incubated for overnight at 37 °C followed by replacement with fresh complete medium and further incubation for the indicated time.

### Real-Time Polymerase Chain Reaction (RT-PCR)

Total cellular RNA was extracted from MELAS cells with a RNeasy Plus Mini Kit (Qiagen, Germantown, MD, USA) according to the manufacturer’s instructions. Complementary DNA (cDNA) was generated by reverse transcriptase polymerase chain reaction (PCR) using the ProtoScript^®^ II First Strand cDNA Synthesis Kit (New England Biolabs, Ipswich, MA, USA). Transcript levels were measured by real time PCR using SYBR green on an ABI 7500 system. Real-time analysis was performed by the ΔΔ^Ct^ method. The primers used were MNRR1 forward: 5′-CACACATGGGTCACGCCATTACT-3′, reverse: 5′-TTCTGGGCACACTCCAGAAACTGT-3′; 18s forward: 5′-CCAGTAAGTGCGGGTCATAA-3′, reverse: 5′-GGCCTCACTAAACCATCCAA-3′; SOD2 forward: 5′-CTGGACAAACCTCAGCCCTA-3′, reverse: 5′-AGCCGTCAGCTTCTCCTTAA-3′; HEY1 forward: 5′-GGCAGGAGGGAAAGGTTACT-3′, reverse: 5′-CTCAGATAACGCGCAACTTC-3′ [27]; TNFα forward: 5′-TGTAGCAAACCCTCAAGCTG -3′, reverse: 5′-GAGGTTGACCTTGGTCTGGT-3′.

### Luciferase Reporter Assay

Luciferase assays were performed with the dual-luciferase reporter assay kit (Promega, Madison, WI, USA). Briefly, cells were lysed in 1x passive lysis buffer (Promega, Madison, WI, USA) and 25 μL of lysate was used for assay with a tube luminometer using an integration time of 10 s. Transfection efficiency was normalized with the co-transfected pRL-SV40 Renilla luciferase expression plasmid [1, 13].

### Immunoblotting

Immunoblotting was performed as described previously [28, 29]. Briefly, cell lysates for immunoblotting were prepared using RIPA buffer (Abcam, Waltham, MA, USA) and included a protease and phosphatase inhibitor cocktail (Sigma-Aldrich, St. Louis, MO, USA). Total protein extracts were obtained by centrifugation at 21,000 × g for 30 min at 4 °C. The clear supernatants were transferred to new tubes and quantified using the Bradford reagent with BSA as standard (BioRad, Hercules, CA, USA). Equal amounts of cell lysates were separated by sodium dodecyl sulfate–polyacrylamide gel electrophoresis (SDS–PAGE), transferred to PVDF membranes (BioRad), and blocked with 5% non-fat dry milk. Incubation with primary antibodies (used at a concentration of 1:500) was performed overnight at 4 °C. The cleaved caspase3 (Catalog # 9661), Histone H3-methylated (H3K4Me3, 9751), PARP (9542), PGC1α (2178), p300 (86377), RBPJk (5313), SIRT1 (2496), GAPDH (8884), actin (12748), and tubulin (9099) antibodies were obtained from Cell Signaling (Danvers, MA, USA). The MNRR1 (19424-1), and SOD2 (24127-1) antibodies were obtained from Proteintech (Chicago, IL). The FLAG antibody (A8592) was obtained from Sigma Aldrich (St. Louis, MO, USA) and the KMT2D antibody (ab213721) was obtained from Abcam (Waltham, MA, USA). Incubation with secondary antibodies (1:5000) was performed for 2 h at room temperature. For detection after immunoblotting, the SuperSignal™ West Pico PLUS substrate or Super Signal™ West Femto Maximum Sensitivity Substrate (ThermoFisher, Waltham, MA, USA) was used to generate chemiluminescence signal, which was detected with X-Ray film (RadTech, Vassar, MI, USA) or BioRad Chemidoc digital imager (Hercules, CA, USA).

### Immunofluorescence

Cells were plated on glass cover slips and treated with MNRR1-FITC conjugated peptide (Peptide 2.0, Chantilly, Virginia, USA) for 48 h, then were fixed with 3.7% formaldehyde (prepared in 1x PBS) at room temperature for 15 min. Cells were washed 3 times with PBST (1x PBS, 0.1% TWEEN-20) for 5 min each and mounted with Vectashield vibrance with DAPI (Cat. # H-1800-10, Vector Labs, Newark, CA, USA). Cells were imaged at 63x on the confocal 60 μm disk setting with the BioTek Cytation C10 using the Gen5 software (version 3.11.19, Agilent, Santa Clara, CA).

### Mitochondria Isolation, blue-native PAGE, and in-gel Complex IV activity

Mitochondria were isolated from scrambled or peptide treated MELAS cells by following previously published protocols [30, 31]. Briefly, cells were pelleted and suspended in Isolation buffer one [225 mM mannitol, 75 mM sucrose, 0.1 mM EGTA, and 30 mM Tris-HCl (pH-7.4)]. Following incubation, homogenization was done by Dounce homogenizer. The homogenates were collected and centrifuged at 600 g for 5 min at 4°C; pellets, containing unbroken cells/tissues and nuclei, were discarded. Supernatants were further centrifuged at 7,000 g for 10 min at 4°C to obtain a mitochondria enriched pellet. The mitochondria containing pellets were washed with Isolation buffer two [225 mM mannitol, 75 mM sucrose, and 30 mM Tris–HCl (pH 7.4)] and centrifuged at 10,000 g for five min at 4°C. The crude mitochondria pellets obtained were resuspended in mitochondria resuspension buffer [250 mM mannitol, 0.5 mM EGTA, and 5 mM HEPES (pH 7.4)]. For BN-PAGE, and for SDS-PAGE, mitochondria were lysed with mitochondria lysis buffer [50 mM Tris-HCl (pH 7.5), 150 mM NaCl, 0.1 mM EDTA, 1% Triton X-100, and 2 mM 6-amino hexanoic acid]. Protein content was quantified by Bradford reagent (Bio-Rad, Hercules, CA, USA).

BN-PAGE was performed on isolated mitochondria by following previously published protocols with minor modifications [32]. Briefly, frozen mitochondria stored at -80°C were thawed on ice and protein was estimated by Bradford reagent. Fifty µg of mitochondria were centrifuged at 10,000 g for 10 min at 4°C; the obtained pellet was solubilized in 20 µL of solubilization buffer (5µL of 4X Native PAGE sample buffer (Invitrogen, Carlsbad, CA, USA), 6 µL of 5% Digitonin (Sigma Aldrich, St. Louis, MO, USA), 9 µL water) and incubated for 20 min on ice. The sample was centrifuged at 20,000 g for 20 min at 4°C and the resultant supernatant was mixed with 1 µL of native loading dye (5% G250 Coomassie Blue in 500 mM 6-aminohexanoic acid). Prepared samples were loaded in a Native PAGE Bis-Tris 3-12% precast gel (Invitrogen) and were run for one hour at 100V in Cathode buffer B followed by replacing it with Cathode buffer B/10 and increasing the voltage to 150V. The entire gel running was performed in a cold room maintaining 4°C. Gels were either stained with Coomassie dye or proceeded for in-gel activity.

In-gel Complex IV activity of mitochondria was visualized on gels following BN-PAGE. Briefly, the gel after BN-PAGE was incubated in a buffer containing 0.5 mg/ml Diaminobenzidine tetrahydrochloride (Sigma), 1 mg/ml Cytochrome *c* (Sigma), and 7.5 mg/ml sucrose in 50 mM phosphate buffer (pH 7.4) at room temperature for about 1 h [33]. The gel was subsequently documented by scanning the image using the BioRad Chemidoc digital imager (Hercules, CA, USA).

### Measurement of Reactive Oxygen Species

Total cellular ROS measurements were performed with CM-H_2_DCFDA (Life Technologies, Carlsbad, CA, USA). Cells were distributed into 96-well plates at 2.5 × 10^4^ cells per well and incubated as described in specific experiments. Cells were then treated with 10 μM CM-H_2_DCFDA in serum-and antibiotic-free medium for 1 h. Cells were washed twice in phosphate-buffered saline and analyzed for fluorescence on a BioTek Synergy H1 Microplate Reader (Agilent, Santa Clara, CA, USA).

### Mitochondrial DNA Levels

Total genomic DNA was isolated from cells expressing each of the mutants using the Invitrogen PureLink Genomic DNA Mini Kit (Thermo Fisher Scientific, Waltham, MA, USA, Catalog # K1820-01) and analyzed by real-time PCR as above. The primer sequences used to amplify mtDNA and GAPDH were as follows: mtDNA forward: 5′-CCTCCCTGTACGAAAGGAC-3′; reverse: 5′-GCGATTAGAATGGGTACAATG-3′; GAPDH forward: 5′-GAGTCAACGGATTTGGTCGT-3′; reverse: 5′-TTGATTTTGGAGGGATCTCG -3′.

### Intact Cellular Oxygen Consumption

Cellular oxygen consumption was measured with a Seahorse XF^e^24 Bioanalyzer (Agilent, Santa Clara, CA, USA). Cells were plated at a concentration of 2 × 10^4^ (MELAS cybrids) per well a day prior to treatment and basal oxygen consumption was measured 48 h after treatments, as described.

### Statistical Analysis

All statistical analyses were performed with the two-sided Wilcoxon rank sum test using MSTAT version 6.1.1 (N. Drinkwater, University of Wisconsin–Madison) or the two-sided t-test (Microsoft Excel). * p< 0.05; ** p < 0.005 *** p < 0.001.

## Results

### Residues 101-113 of MNRR1 are required and sufficient for transcriptional activation of the *MNRR1* promoter

In 143B cells containing 73% of the MELAS m.3243A>G mtDNA mutation (DW7, MELAS cells), MNRR1 expression promotes mitochondrial oxygen consumption and can do so even when it is mutationally excluded from the mitochondria (**Fig. 1A**). Our previous work has identified mitochondrial biogenesis as one of the pathways activated by MNRR1 [19] and could underlie the observed effect. Since MNRR1 carries out multiple functions in at least two organelles [13, 16], we sought to identify the minimal domain necessary for transcriptional activation. To do so, we generated MNRR1 deletion mutants (**Fig. 1B**) and confirmed their ability to be expressed in the MELAS cells (**Fig. 1C**). Since MNRR1 activates its own transcription [1, 16], we used the 952-bp *MNRR1-*promoter reporter that drives expression of luciferase to measure activity [1, 16] and found that residues 101-113 (**Fig. 1B, construct #5, 1D**) of MNRR1 protein were required for transcriptional activation. We then used a known downstream transcriptional target [1], PGC1α (Peroxisome proliferator-activated receptor Gamma Coactivator-1alpha), as a readout to confirm MNRR1 transcriptional activation. Overexpressing the TA-domain harboring protein (**Fig. 1B, construct #5**) in MELAS cells increased the levels of PGC1α over that observed in cells expressing the full-length wild-type protein (WT) (**Fig. 1E),** indicating that the TA-domain is sufficient for rescue of the phenotype.

**Figure 1:**
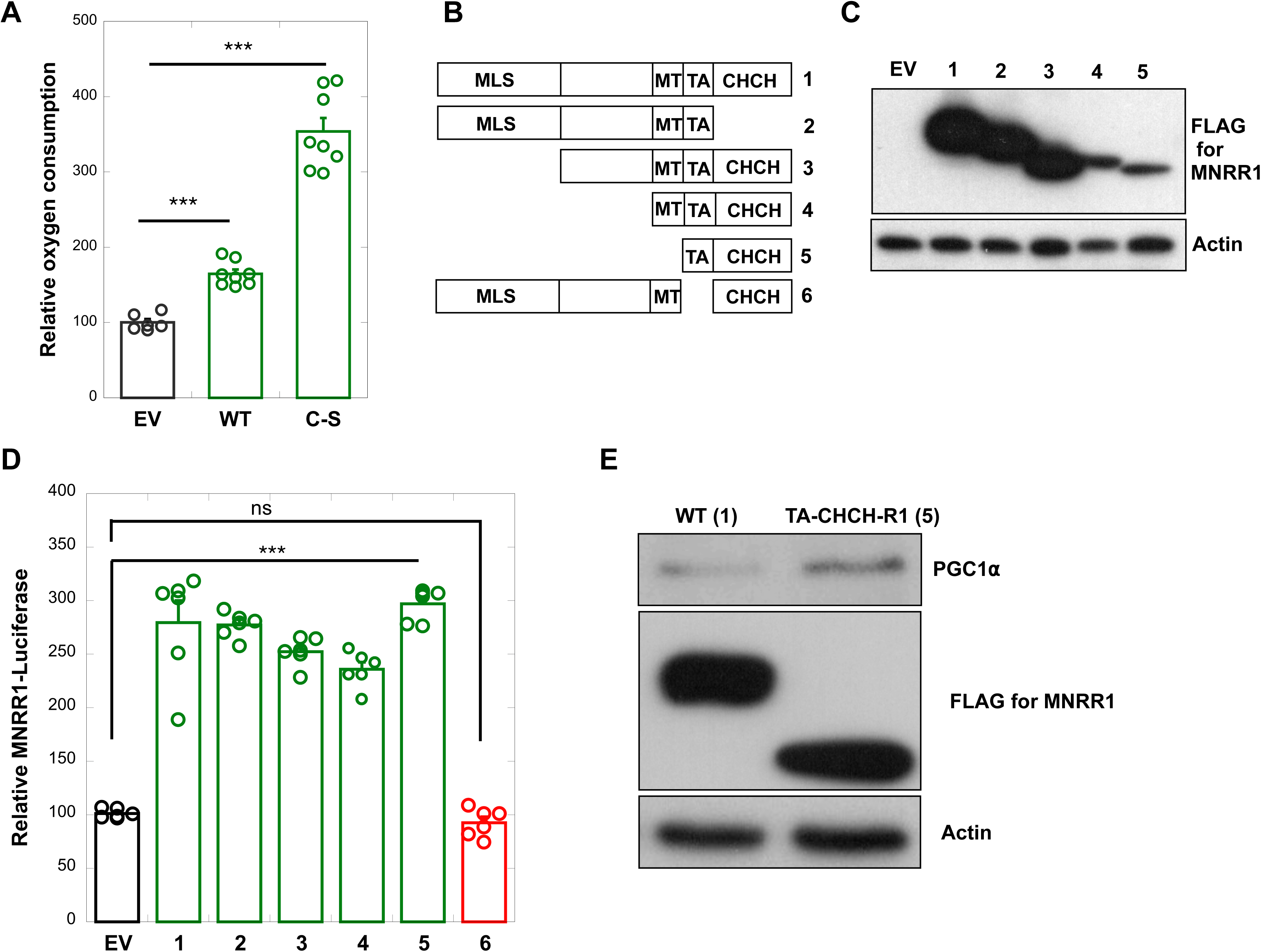
Amino acids 101-113 are required for transcriptional activation function of MNRR1. **A)** Basal mitochondrial oxygen consumption in DW7 MELAS cells expressing either an empty vector (EV), wild-type MNRR1 (WT), or the nuclear localized MNRR1 (C-S, cysteine residues mutated to serine). ***p<0.001. **B)** Depiction of Flag-tagged MNRR1 protein domain deletion mutants used in the study. MLS: Mitochondrial Localization Sequence, MT: Mitochondrial domain required for interaction with COX, TA: Transcriptional activation domain, and CHCH: Coiled-coil helix, coiled-coil helix domain. **C)** Immunoblotting of cell lysates expressing each of the deletion mutants in DW7 MELAS cells. Actin used as a loading control. **D)** Activation of MNRR1-promoter luciferase by each of the MNRR1 mutants (1 through 6) relative to that of the empty vector (EV). ***p<0.001. **E)** MNRR1 construct # 5 (Fig.1B) expressing the TA domain along with the CHCH domain (TA-CHCH-R1) transfected in DW7 cells display increase in the protein levels of PGC1α, a downstream transcriptional target of MNRR1 when compared to cells transfected with wild type (WT, construct #1,Fig 1B). Expression levels of WT and mutant protein shown. Actin used as a loading control.

### The TA-R1-peptide can carry out MNRR1’s nuclear functions

Having shown that MNRR1 protein residues 101-113 could promote transcriptional activation of *MNRR1* containing the TA domain (**Fig. 1D**) and could stimulate one of its downstream protein targets, PGC1⍺ (**Fig. 1E**), we generated a 20 amino acid peptide (TA-R1) encompassing the transcriptional domain of MNRR1 and examined its effects on nuclear function in more detail **(Supp. Fig. 1A)**. We first determined that the FITC-tagged TA-R1 peptide could localize to the nucleus using immunofluorescent staining **(Supp. Fig. 1B)**. Each of the peptides (TA-R1, D1, D2, and D3) were able to activate the MNRR1-promoter luciferase construct **(Supp. Fig. 1C)** and MNRR1 protein levels **(Supp. Fig. 1D).** Treatment with the TA-peptide is able to increase transcription of MNRR1 in proportion to amount of peptide added (**Fig. 2A**), an effect also seen at the protein level (**Fig. 2B**). Furthermore, TA-R1 peptide increased the amounts of two downstream targets, PGC1α and SOD, both at the level of transcription (**Fig. 2C**) and translation (**Fig. 2D**), in a concentration dependent manner. The peptide-stimulated increase in MNRR1 expression is not restricted to the MELAS cells but also takes place in a variety of wild-type as well as disease carrying human cell lines (**Supp. Fig. 2A).** MNRR1-responsive genes (PGC1α and SOD) harbor a common promoter element needed for MNRR1 transcriptional activation and response to hypoxia [1, 16], termed the oxygen responsive element (ORE). We previously showed that the ORE, which is a consensus binding site for RBPJκ (Recombination Signal Binding Protein for Immunoglobulin Kappa J Region) [13], functions towards transcriptional activation by MNRR1 by indeed binding the transcription factor RBPJκ. The TA-R1 peptide functions only at the ORE as the peptide fails to activate a reporter with no promoter (basal reporter) (**Supp. Fig. 2B**) or a specific deletion of the ORE (Δ ORE) (**Supp. Fig. 2C**). Furthermore, the peptide also increases transcription of the reporter construct in Rho^0^ cells that lack functional mitochondrial DNA (**Fig. 2E**). Taken together, these results suggest that the TA-R1 peptide predominantly localizes to the nucleus and functions as a transcriptional regulator at the ORE, similarly to the full-length, wild-type MNRR1. The TA domain of MNRR1 is important as a mutation in this domain (Q112H) has been associated with CMT1A [29] and results in defective transcriptional activation. This data indicates that amino acids 109-113 are required for and play a vital role in transcriptional activation by MNRR1.

**Figure 2:**
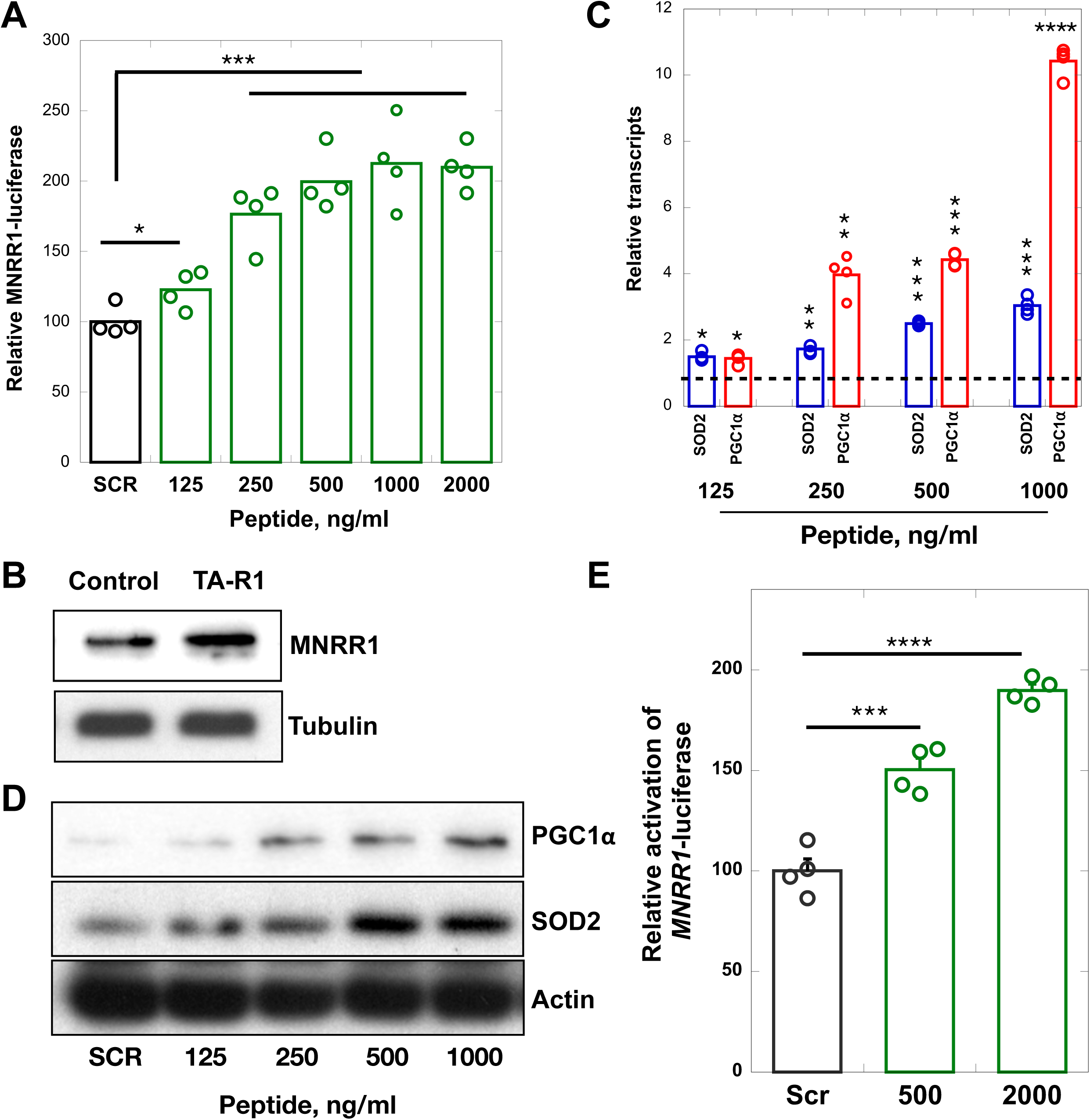
MNRR1 peptide activates transcription of downstream genes. **A)** MNRR1 peptide treated cells increased activity of the MNRR1-promoter luciferase in a concentration dependent manner relative to that of cells treated with a scrambled peptide (2000 ng/ml). *p<0.05, ***p<0.001. **B)** Western blot of lysates from MNRR1 peptide (TA-R1) treated DW7 MELAS cells showing an increase in the levels of MNRR1 protein. Tubulin used as a loading control. TA-R1 increases the transcript (**C)** and protein **(D)** levels of *SOD2* and *PGC1α* in a concentration dependent manner relative to the levels observed with the scrambled peptide, depicted in **(C**) by a dotted line. *p<0.05. **E)** MNRR1 peptide also increases luciferase reporter activity in Rho^0^ cells. *p<0.05.

### MNRR1 activates transcription via recruitment of the histone acetyltransferase p300 to the RBPJ**_κ_** promoter complex

The mechanism by which RBPJ_κ_ regulates transcription has been partially characterized [34]. Briefly, during transcriptional repression, the repressor proteins SHARP (SMRT/HDAC1 Associated Repressor Protein) and HDAC1 are bound to RBPJ_κ_ and recruit the nuclear co-repressor complex (nCOR). During activatory priming, nCOR is replaced by Histone-lysine N-methyltransferase 2D (KMT2D), the enzyme responsible for increasing levels of H3K4me3 [35], a histone signature associated with active promoters [36]. Further, the KMT2D-SHARP complex is displaced to recruit the co-activator p300 (**Fig. 3A**). We therefore hypothesized that MNRR1 functions by facilitating recruitment of p300 to RBPJ_κ_. We initially tested this hypothesis by co-immunoprecipitation and could show enhanced interaction between RBPJ_κ_ and p300 in the presence of MNRR1 and a relative decrease of CXXC5 (**Fig. 3B)**. MNRR1 expressing cells also displayed an increase in the levels of H3K4Me3 (**Fig. 3C**). The C-terminus of the protein encompassing the TA domain is sufficient for this increase (**Fig. 3D**). In the presence of MNRR1, the interaction between RBPJ_κ_ and H3K4me3-modified histones is increased (**Fig. 3E**), suggestive of the transcriptional activatory role of MNRR1. Co-expression of MNRR1 along with Sirt1 (the deacetylase for residue K119 on MNRR1, which enhances its transcriptional activation function) shows recruitment of p300 with a concomitant displacement of KMT2D (**Fig. 3F**). In support of the role of the deacetylated mutant of MNRR1 (K-R) in the recruitment of p300, the acetylmimetic mutant (K-Q) facilitates interaction of RBPJ_κ_ with KMT2D (**Fig. 3G**). These results suggest that removal of acetylation is a potential regulatory step in promoter activation. To then test if the TA-R1 peptide is sufficient to recruit p300 to RBPJ_κ_, we performed immunoprecipitation of RBPJ_κ_ in the presence of TA-R1. As shown in **Fig. 4A**, TA-R1 facilitates the interaction between p300 and RBPJ_κ_. This peptide mediated interaction is also facilitated in MNRR1-knockout cells (**Fig. 4B**), suggesting that the minimal peptide is sufficient to activate transcription at the ORE. Finally, the MNRR1 peptide induces transcription of a bonafide RBPJ_κ_ target, *Hey1* **(Fig. 4C)** [27]. These results indicate that MNRR1 mechanistically functions as a transcriptional co-activator of RBPJκ by recruiting the histone acetyl transferase p300 to the transcriptional site.

**Figure 3:**
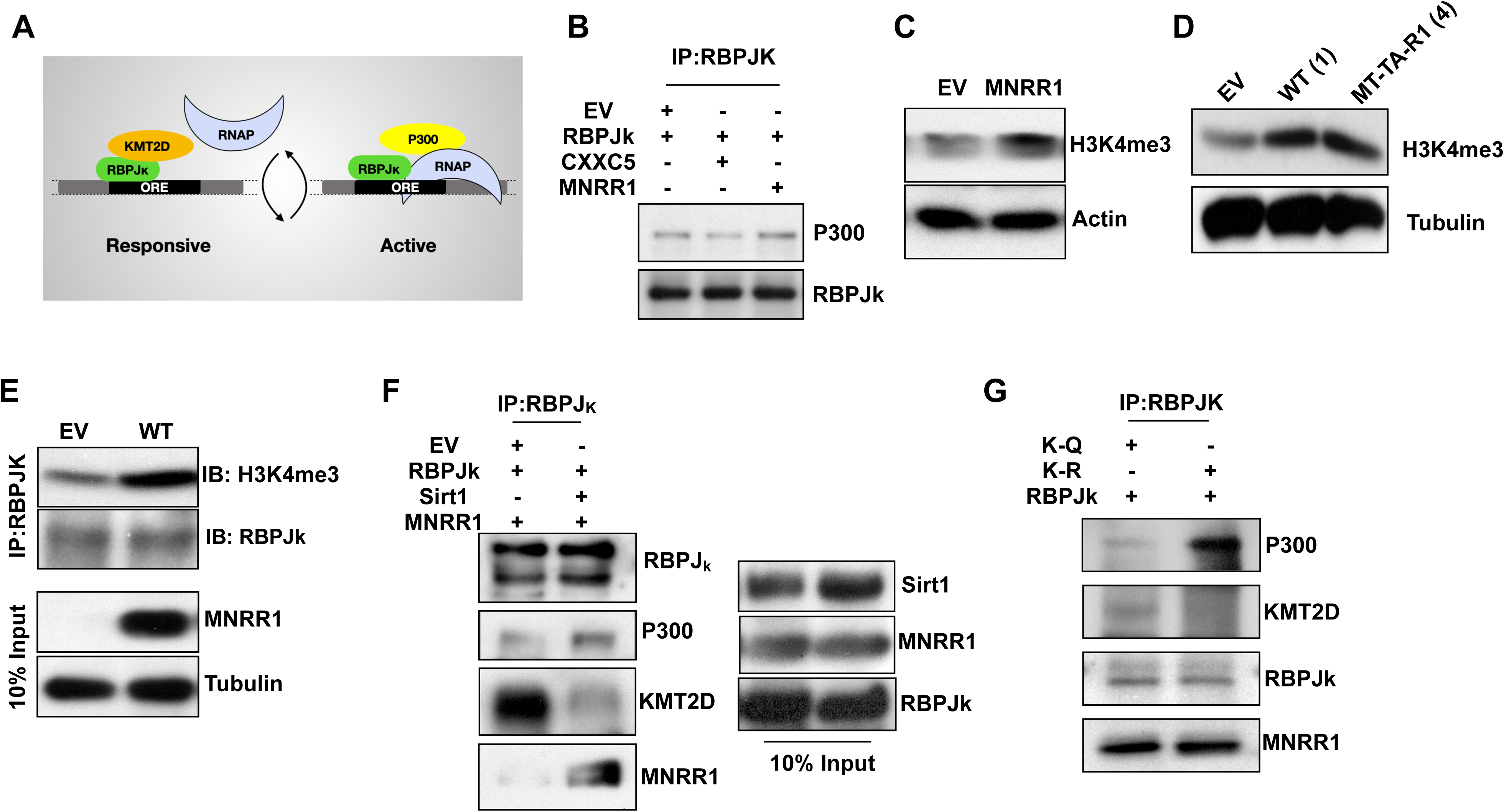
MNRR1 recruits p300 to RBPJκ at the ORE. **A)** Model for activation of transcription at the ORE by RBPJκ (adapted from [34]). RNAP, RNA polymerase; KMT2D, Lysine Methyltransferase 2D. **B)** Co-immunoprecipitation analysis of RBPJκ and p300 in cells expressing the indicated plasmids. **C)** MNRR1 overexpressing cells display an increase in the levels of H3K4me3, a marker of active transcription. **D)** MT-TA-CHCH (construct # 4, Fig.1B) is sufficient to induce the activatory histone signature H3K4me3. **E)** Cells expressing wild type MNRR1 (WT) display an increased interaction between RBPJκ and H3K4me3 compared to cells expressing an empty vector (EV). **F)** MNRR1 in the presence of Sirt1 (that deacetylates it, thereby making it a superior transcription factor) displaces KMT2D and recruits p300 to RBPJκ. **G)** Cells expressing the deacetylated (K-R) but not the acetylmimetic (K-Q) mutant of MNRR1 display enhanced interaction between RBPJκ and p300.

**Figure 4:**
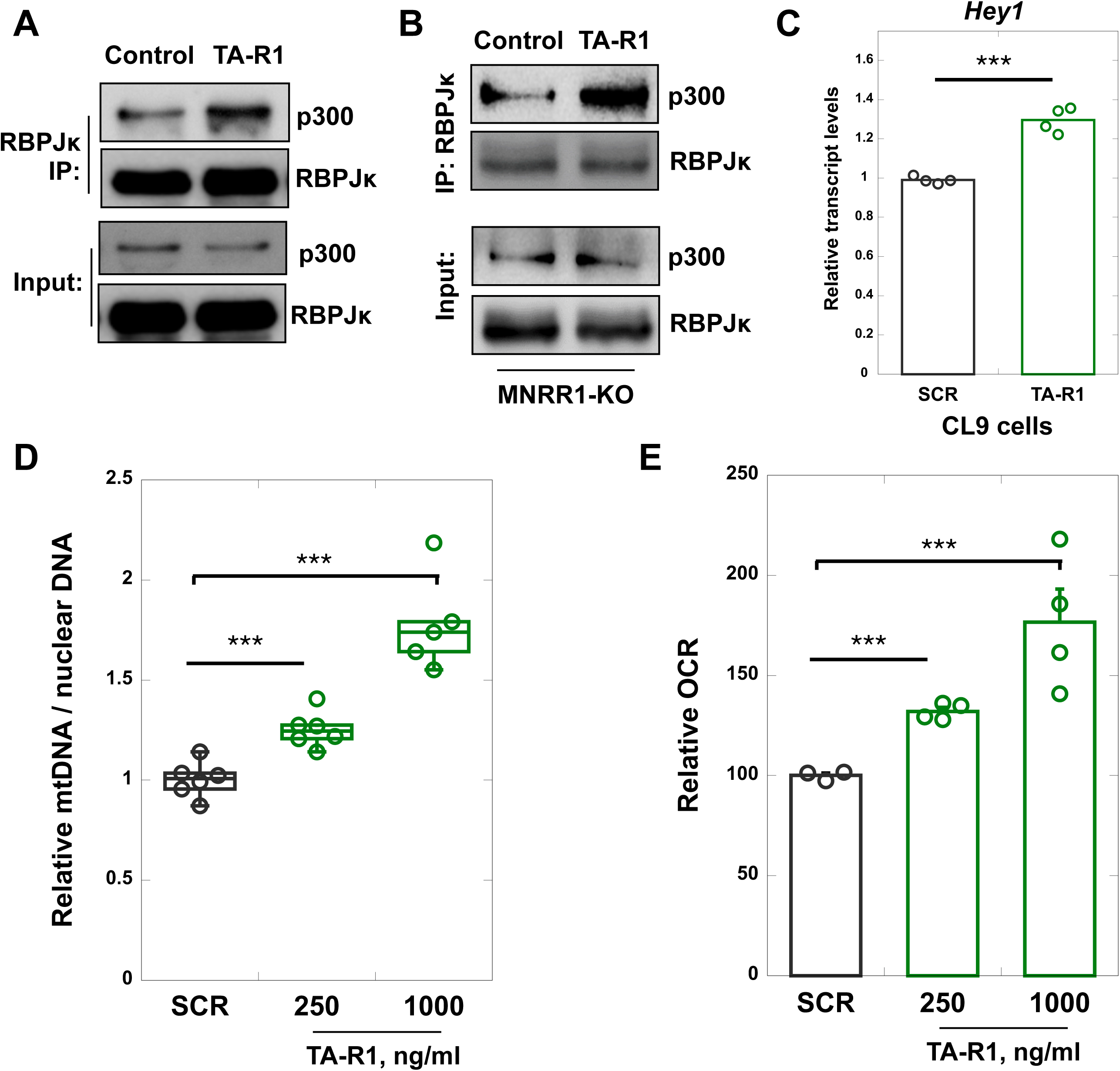
MNRR1 peptide recruits p300 to RBPJκ. **A)** Immunoprecipitation showing TA-R1 treated MELAS (1000 ng/mL for 48h) cells display enhanced interaction between RBPJκ and p300. **B)** TA-R1 also recruits p300 in the absence of endogenous MNRR1 (MNRR1-knockout, MDAMB231-MNRR1-KO) cells treated for 48 h (1000 ng/mL). **C)** TA-R1 increases the transcript levels of *Hey1*, a bonafide RBPJκ transcriptional target in MELAS cells (treated with 1000 ng/mL peptide for 48h).***p<0.001. **D)** TA-R1 treated DW7 cells (1000 ng/mL peptide for 48h) have increased mitochondrial DNA levels (mtDNA). ***p<0.001. **E)** TA-R1 increases basal oxygen consumption rate in DW7 cells (treated with 1000 ng/mL peptide for 48h). ***p<0.001.

### Mitochondrial effects of the TA-R1-peptide

Since MNRR1 is a regulator of mitochondrial function, we examined the effects of the peptide on mitochondrial genes that are transcriptionally regulated by MNRR1 such as *PGC1α* [1] and the mitochondrial antioxidant *SOD2* [16] and found these genes to be transcriptionally stimulated (**Fig. 2C**). We confirmed their upregulation at the protein level as well (**Fig. 2D**). Next, we confirmed the functional consequences of their upregulation by assessing the levels of mitochondrial DNA (**Fig. 4D**) and of mitochondrial respiration (**Fig. 4E**) and found these to be enhanced as expected. Two direct effects of MNRR1 in the mitochondria that can be differentiated on the basis of Y99 phosphorylation are ATP production and resistance to apoptosis. To further validate the specificity of the peptide to function exclusively in the nucleus, we tested for the effects of the peptide mitochondrial specific processes. Although MNRR1 is known to be able to block apoptosis [18], cells treated with the TA-R1 peptide displayed similar levels of the apoptotic markers PARP and Caspase 3 (**Supp. Fig. 2D).** In addition, although MNRR1 expression stimulates complex IV activity [16], the peptide did not affect in-gel Complex IV activity (**Supp. Fig. 2E**), suggesting that the effects on mitochondrial function are probably via nuclear function of MNRR1. To assess the effects on nucleus specific processes, we have previously shown that MNRR1 nuclear function can rescue Lipopolysaccharide (LPS)-induced TNFα expression and reactive oxygen species (ROS) levels [7, 8]. TA-peptide treated MELAS cells displayed a significant reduction in LPS-induced inflammatory response (**Supp. Fig. 2F)** as well as a reduction in ROS levels (**Supp. Fig. 2G)**.

## Conclusions

a. MNRR1 functions in the nucleus by recruiting the histone acetyl transferase p300.
b. The transcriptional domain of MNRR1 is necessary and sufficient for nuclear function.
c. MNRR1 peptides can induce downstream homeostatic genes in a cellular model of MELAS.
d. MNRR1 peptides are promising therapeutically to activate mitochondrial function.

## Author Contributions

N.P, V.P, D.P, and SR performed the experiments. N.P, L.I.G, and S.A participated in experimental design, data analysis, and writing the manuscript. All authors have read and agreed to the published version of the manuscript.

## Data Availability Statement

All study data presented in this manuscript are included in the article or are available from the lead contacts upon request. Unique reagents generated from this study are available from the lead contacts with a completed Materials Transfer Agreement. This study did not generate original code.

---

**Supplementary Figure 1: A)** MNRR1 peptide sequences used in the study. **B)** Immunofluorescence microscopy of FITC-tagged TA-R1 peptide showing an overlap with DAPI for nuclear localization MELAS cells were treated with 1000 ng/mL TA-R1 peptide added to the medium for 48 h. **C)** *MNRR1*-luciferase reporter activity of the individual peptides added to growth medium at 1000 ng/mL for 48 h. **p<0.005. **D)** Endogenous MNRR1 protein levels increase when cells are treated with minimal MNRR1 peptides. Cells were treated with the indicated peptides at 1000 ng/mL for 48 h. Tubulin used as a loading control.

**Supplementary Figure 2: A)** TA-R1 peptide increases endogenous MNRR1 protein levels in the indicated cell lines. **B)** TA-R1 does not activate the basal (promoter-less, pGL4-basic) luciferase reporter. **C)** TA-R1 does not activate the MNRR1 reporter harboring a deletion of the ORE (ΔORE). **D)** TA-R1 treated cells do not display any changes in apoptotic marker PARP and cleaved caspase 3. **E)** TA-R1 cells do not display a change in complex IV activity on a blue native gel. Activity was measured with diaminobenzidine. **F)** TA-R1 rescues the LPS-induced increase in the levels of an inflammatory marker, TNFα. **p<0.005. Co-treatment with LPS (0.5 µg/mL) and TA-R1 (1000 ng/ mL) was done for 48h. **G)** TA-R1 also reduces ROS levels that are increased upon treatment with LPS. **p<0.005. Conditions were as in (F). In experiments shown in (A) – (E), TA-R1 was added at 1000 ng/mL for 48 h.

