## Supplementary Data for "Transcriptional activation by MNRR1 is effected by recruiting p300 and can be induced by minimal peptides"

**A**

| Peptide | Sequence |
|---|---|
| TA-R1 | EPQGTQPAQQQQPCLY |
| D1 | AQQQQPCLYEIKQ |
| D2 | QPAQQQQP |
| D3 | EPQGTQPAQQQQPCL |

**B**

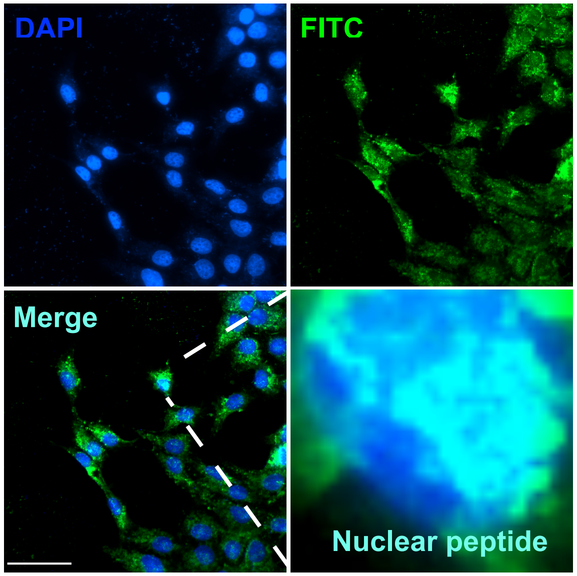


**C**

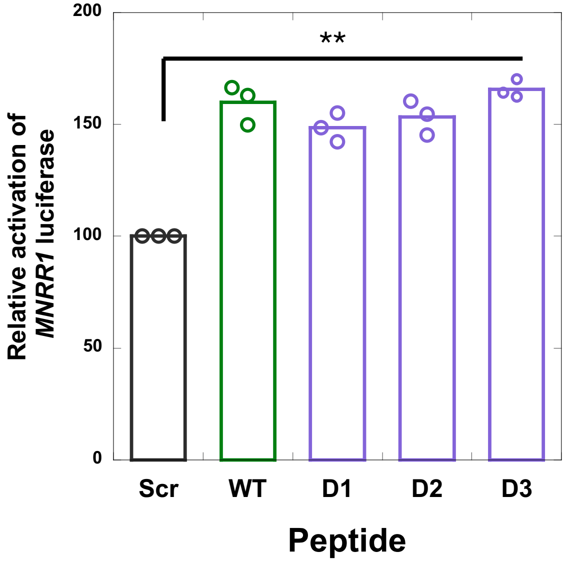


**D**

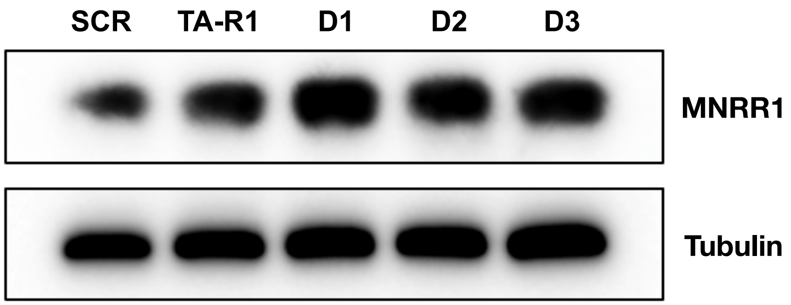


### Supplementary Figure 1

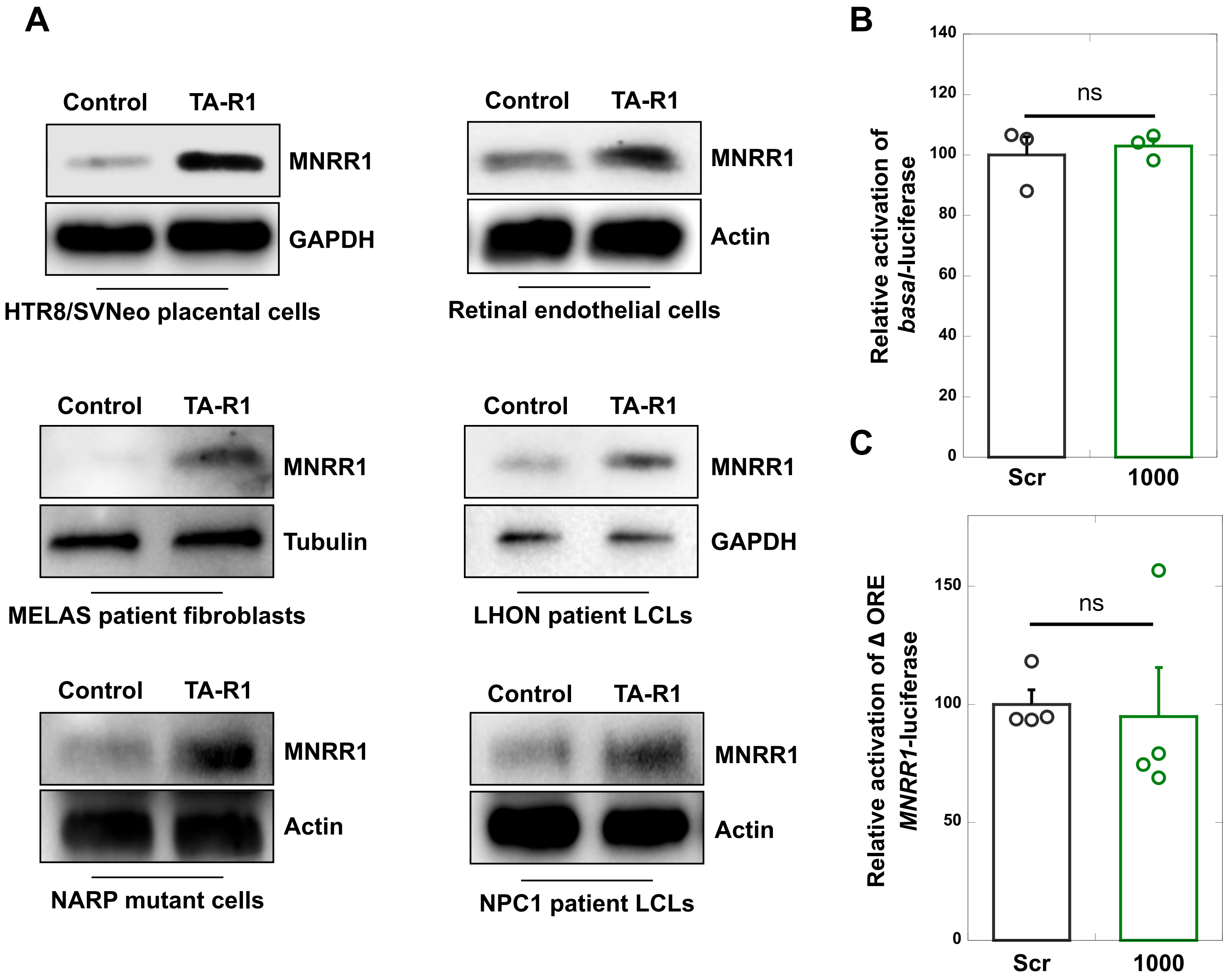


### Supplementary Figure 2

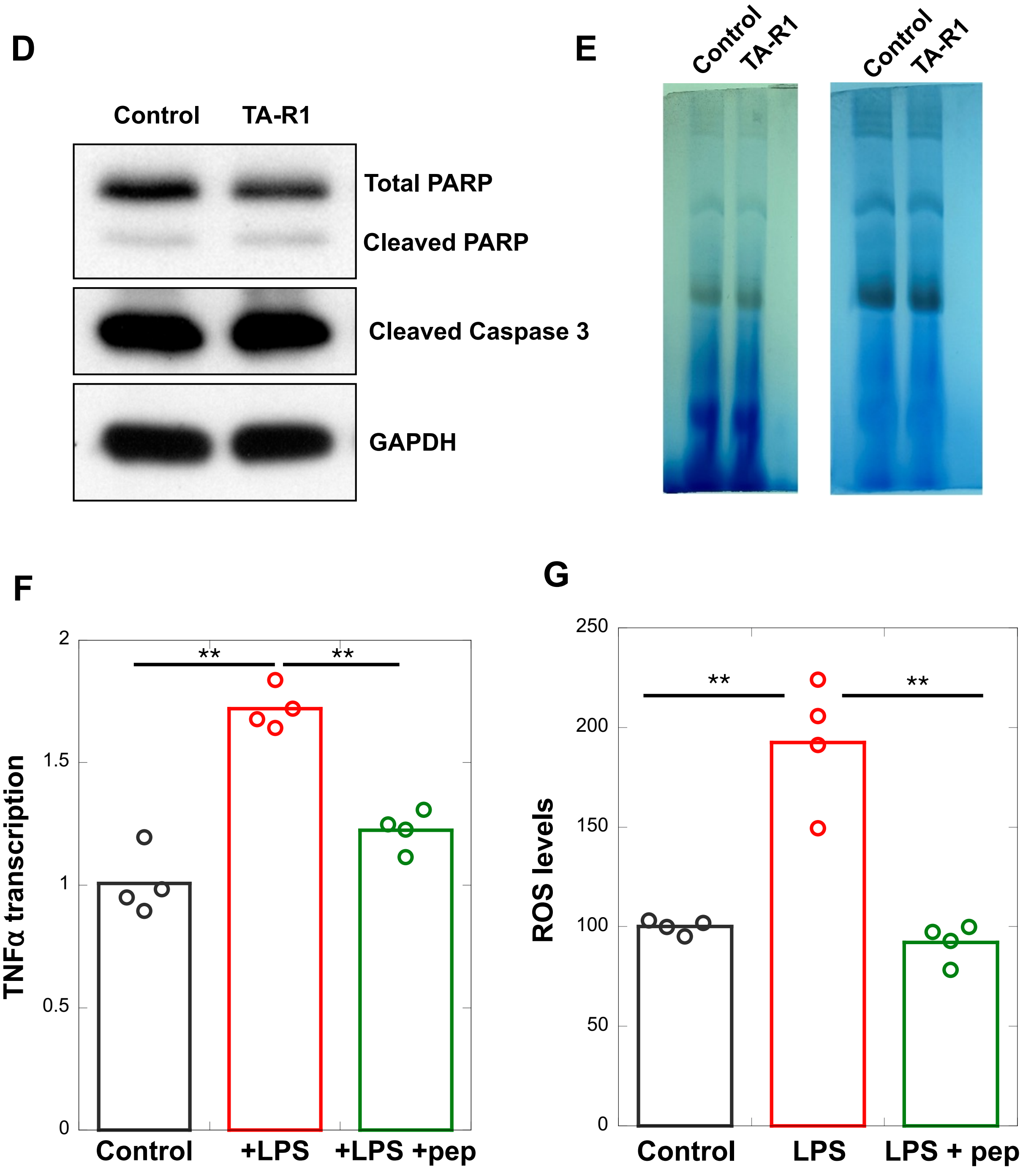


### Supplementary Figure 2
